# Pulsed inhibition of corticospinal excitability by the thalamocortical sleep spindle

**DOI:** 10.1101/2024.07.22.604668

**Authors:** Umair Hassan, Prince Okyere, Milad Amini Masouleh, Christoph Zrenner, Ulf Ziemann, Til Ole Bergmann

## Abstract

Thalamocortical sleep spindles, i.e., oscillatory bursts at ∼12-15 Hz of waxing and waning amplitude, are a hallmark feature of non-rapid eye movement (NREM) sleep and believed to play a key role in sleep-dependent memory reactivation and consolidation. Generated in the thalamus and projecting to neocortex and hippocampus, they are phasically modulated by neocortical slow oscillations (<1 Hz) and in turn phasically modulate hippocampal sharp-wave ripples (>80 Hz). This hierarchical cross-frequency nesting may enable phase-dependent plasticity in the neocortex, and spindles have thus been considered windows of plasticity in the sleeping brain. However, the assumed phasic excitability modulation had not yet been demonstrated for spindles. Utilizing a recently developed real-time spindle detection algorithm, we applied spindle phase-triggered transcranial magnetic stimulation (TMS) to the primary motor cortex (M1) hand area and measured motor evoked potentials (MEP) to characterize corticospinal excitability during sleep spindles. We found a net suppression of MEP amplitudes during spindles, driven by selective inhibition during the falling flank of the spindle oscillation, but no inhibition during its peak, rising flank, and trough. Importantly, this phasic inhibition occurred on top of the general sleep-related inhibition observed during spindle-free NREM sleep and did not extend into the immediate refractory post-spindle periods. We conclude that spindles exert asymmetric “pulsed inhibition” of corticospinal excitability, which is assumedly relevant for processes of phase-dependent plasticity. These findings and the developed real-time spindle targeting methods will enable future studies to uncover the causal role of spindles in synaptic plasticity and systems memory consolidation.

## 1 Introduction

Thalamocortical sleep spindles are transient oscillatory complexes of waxing and waning amplitude at sigma frequency (∼12-15 Hz) and constitute a hallmark feature of the electroencephalogram (EEG) during non-rapid eye movement (NREM) sleep. Nested phase–amplitude coupling of spindles with 0.5-1 Hz neocortical slow oscillations (SOs) and hippocampal ripples (> 80 Hz) (Staresina et al. 2015) facilitates sleep-dependent memory reactivation (Bergmann et al. 2012) and systems consolidation (Diekelmann and Born 2010), thus endorsing the concepts of phase-dependent plasticity (Bergmann and Born 2018) and synaptic rescaling of cortical neurons (Klinzing et al. 2019). Sleep spindles are generated by the reciprocal interaction of inhibitory reticular-thalamic and excitatory corticothalamic neurons (Fernandez and Lüthi 2020), while their initiation and termination may involve both excitatory corticothalamic (Bal and McCormick 1996; Bonjean et al. 2011; Destexhe et al. 1996; Lüthi and McCormick 1998; Lüthi and McCormick 1999; Timofeev 2001) and inhibitory reticular-thalamic neurons (Barthó et al. 2014; Langdon et al. 2012; Rovo et al. 2014). These mechanisms of spindle generation and termination likely cause rhythmic fluctuations in the excitation/inhibition balance (EIB) of neocortical neuron populations (Brécier et al. 2022; Niethard et al. 2018; Peyrache et al. 2011). Furthermore, stimulation of cortical pyramidal cells with spindle-like 12-15 Hz pulse sequences promoted Ca^2+^-dependent long-term potentiation (LTP) of excitatory postsynaptic potentials, suggesting that spindles may also contribute to spike timing-dependent plasticity (STDP) via Ca^2+^ influx into cortical neurons (Dickey et al. 2021; Feldman 2012; Rosanova 2005). The hypothesis of ’phase-dependent plasticity’ assumes particularly strong Ca^2+^ influx and associated LTP when spindles are precisely timed to the depolarizing phase of neocortical SOs (Bergmann and Born 2018; Helfrich et al. 2018) and ripples to the most excitable phase of spindles (Staresina et al. 2015). Notably, the same neurophysiological mechanisms underlying EIB fluctuations during spindles and SOs are presumably also responsible for the synaptic rescaling of neocortical synapses as a mechanism of systems memory consolidation during sleep (Klinzing et al. 2019; Niethard et al. 2016; Niethard et al. 2017; Niethard et al. 2021). However, while rapid neocortical excitability fluctuations during the SO had already been demonstrated in humans (Bergmann et al. 2012), a similar phasic modulation had yet to be shown for sleep spindles. Here, we conducted a first investigation into the cortical excitability dynamics during sleep spindles in humans by translating the approach of real-time EEG phase triggered transcranial magnetic stimulation (TMS) and motor evoked potential (MEP) measurements previously used to study excitability dynamics of the sensorimotor mu-alpha rhythm (Bergmann et al. 2019; Zrenner et al. 2018) during wakefulness and the SO during sleep (Bergmann et al. 2012). Specifically, we used a recently developed real-time spindle detection algorithm (Hassan et al. 2022) to trigger TMS in a spindle phase-specific manner and assess corticospinal excitability, as indexed by the MEP amplitude in response to a single TMS pulse delivered over left primary motor cortex (M1) hand area. We probed excitability at four different phase angles (trough, rising flank, peak, falling flank) close to the center of spontaneously occurring spindles, as well as during the refractory period immediately following spindles, and during baseline periods of desynchronized low amplitude NREM sleep where spindles and SO were absent (**Figure 1B**). The derived cortical excitability profile of the sleep spindle shows a pattern of asymmetric pulsed inhibition, with phasic inhibition only being evident during the falling flanks of the oscillation.

**Figure 1:**
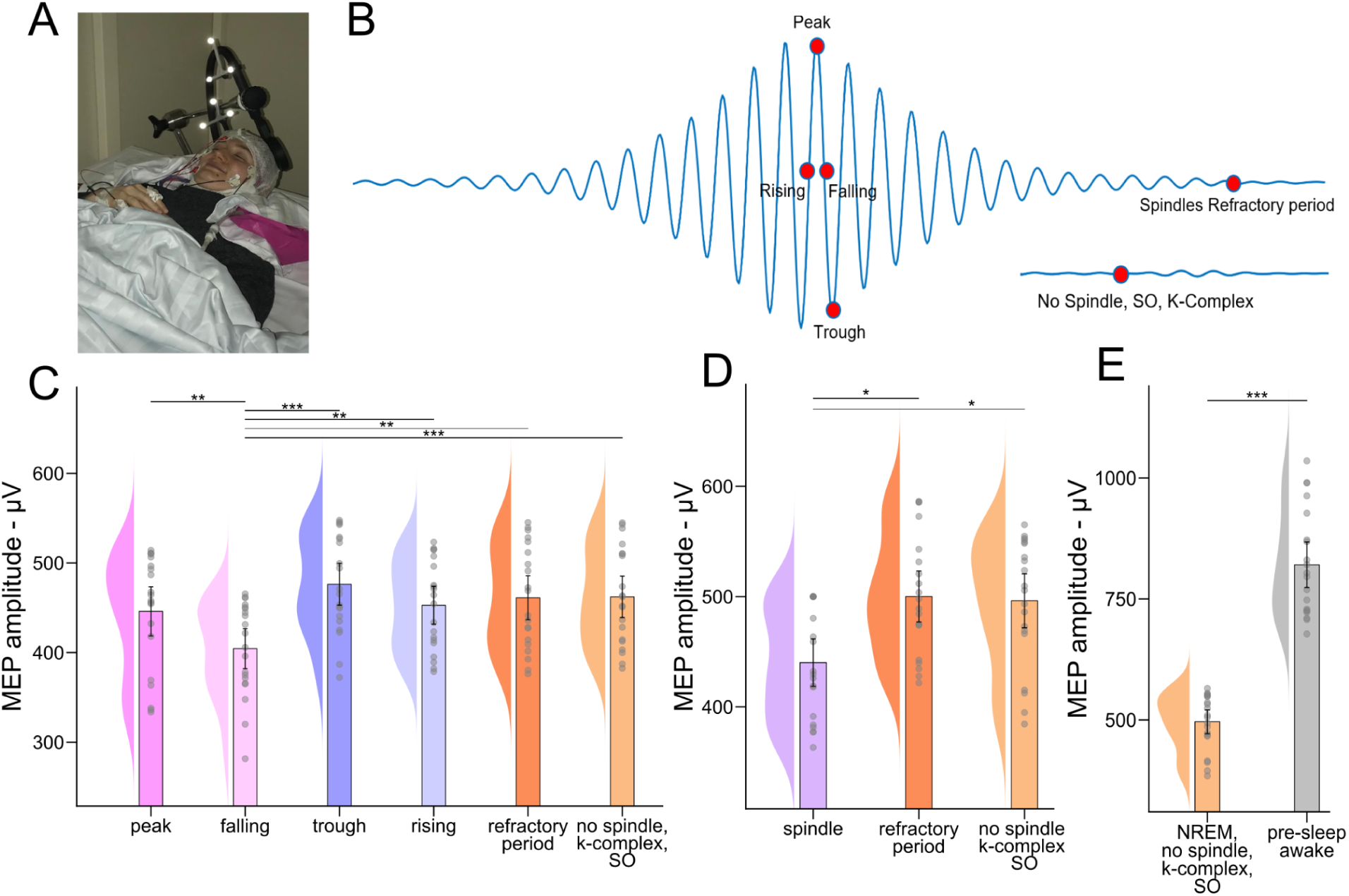
Real-time EEG spindle phase-triggered TMS. **(A)** Experimental setup. **(B)** Four distinct phases of the spindle oscillation (trough, rising flank, peak, falling flank) were targeted by single-pulse TMS and compared to spindle- and SO-free intervals and immediately following (post) sleep spindles during refractory period.Significance of post hoc comparisons is indicated as follows: *p < 0.05; **p < 0.01; ; ***p < 0.001. **(C)** During spindles, MEP amplitudes revealed a rhythmic inhibition of motor corticospinal excitability only at the falling flank. **(D)** Averaged across all spindle phases, the MEP amplitude is still smaller than during spindle-free and spindle-refractory periods. **(E)** During spindle-free baseline NREM periods, MEP amplitude is reduced relative to pre-sleep wakefulness.

## 2 Materials and Methods

### 2.1 Subjects

Twenty (N = 20) healthy, right-handed volunteers (12 females; mean age, 23; range, 18–42 years), who were free of medication, had no history of neurological or psychiatric disease, nor any contraindication to TMS (Rossi et al. 2011), participated after providing written informed consent. They were not permitted to consume alcohol or caffeine on the day of the experiment, and they were required to be awake for at least 8 hours prior to the experiment to maintain moderate sleep pressure. The study protocol followed the Declaration of Helsinki and was approved by the local ethics committee of the Landesärztekammer Rheinland-Pfalz. Subjects were recruited based on the following inclusion criteria: (1) ability to sleep on their back in supine position for at least an hour without strong snoring (head movements) under the inconvenient experimental conditions; and (2) the presence of a TMS motor hotspot for evoking consistent MEPs of sufficient amplitude (with scalp-cortex distance increased by the EEG cap, reduced stimulator output due to the extra-long coil cable, and the know decrease in MEP amplitudes during sleep) in either the right first dorsal interosseous (FDI) or abductor pollicis brevis (APB) hand muscles. In total, 20 of 31 screened subjects met these criteria, were recruited, and completed the study.

### 2.2 Experimental design & general procedure

Subjects participated in two nocturnal nap sessions, the adaptation session and the experimental session, which included several preparatory measures as well as the main experiment. Preparatory measures during both sessions (see below for details) included: placement of EEG, polysomnography (PSG), and hand muscle electromyography (EMG) electrodes, arrangements for TMS neuronavigation, motor hotspot search, automated determination of resting motor threshold (RMT), and MEP measurements during wakefulness pre-sleep. Participants arrived in the laboratory after at least 8 hours of wakefulness during both sessions (Time of arrival (hours - HH), Mean: 19, SD: 04, Range: 14-02). After pre-sleep TMS measurements, they were allowed to fall asleep, sleep for the first few sleep cycles, and then leave the laboratory. Subjects slept in a soundproof and electromagnetically shielded sleeping cabin (Desone, Germany) in a wooden bed on a comfortable mattress, lying on their back in supine position. They were covered with a blanket and had their heads supported by a vacuum cushion. Single-pulse TMS was delivered to the left M1 hand area during the experimental nap session to assess corticospinal excitability. TMS was automatically triggered in real-time (see below for details) to target six different NREM sleep periods: four different phase angles during sleep spindles, i.e., either (1) the peak (0°), (2) the falling flank (90°), (3) the trough (180°), (4) the rising flank (270°), (5) the spindle refractory period (i.e., 1.5 s after the spindle center), or (6) desynchronized periods of NREM sleep where spindles, SOs, or K-complexes were absent (collectively referred as spindle-free condition) (**Figure 1B**). The order of these six different experimental conditions was pseudo-randomized across trials to avoid any systematic carry-over effects between conditions.

### 2.3 EEG, PSG & EMG recordings

64-channel EEG was recorded using a textile EEG cap with extra-flat TMS-compatible sintered Ag/AgCl electrodes (Multitrodes-TMS, EasyCap; channels Fp1, Fp2, Fpz, AF7, AF3, AFz, AF4, AF8, F9, F7, F5, F3, F1, Fz, F2, F4, F6, F8, F10, FT9, FT7, FC5, FC3, FC1, FC2, FC4, FC6, FT8, FT10, T7, C5, C3, C1, Cz, C2, C4, C6, T8, TP7, CP5, CP3, CP1, CPz, CP2, CP4, CP6, TP8, P7, P5, P3, P1, Pz, P2, P4, P6, P8, PO7, PO3, PO4, PO8, O1, O2, M1 (left mastoid), M2 (right mastoid); Reference, FCz; Ground, POz). For PSG, EMG at the chin, and the vertical and horizontal electrooculogram (VEOG and HEOG) were recorded using bipolar electrode montages. Surface EMG was recorded with disposable gel electrodes from the right FDI, APB, and ADM muscles in a belly-tendon montage. EEG, PSG, and EMG were digitized in DC mode at 5-kHz sampling rate and with 1250-Hz anti-aliasing low-pass filter, using a TMS-compatible 24-bit amplifier (NeurOne Tesla with Digital-Out Option, Bittium, Finland) connected to an 8-V battery.

### 2.4 TMS

TMS was applied to the left M1 hand region via figure-of-eight C-B60 coil with an outer diameter of 75 mm connected to a MagPro-X100 stimulator (MagVenture, Denmark). The coil was placed on the head from anteromedial (coil handle) to posterolateral (coil head), and a biphasic pulse with a reversed current direction induced posterolateral-to-anteromedial current in brain tissue for the second, more effective, half-wave. The MagVenture FlexArm® attached to the base of the wooden bed in the sleep lab held the TMS coil. The coil position producing consistent MEPs in the right FDI or APB hand muscle (i.e., the ‘hotspot’) was saved and maintained using MR-template-based frameless stereotactic neuronavigation (Localite, Germany). The BEST toolbox (Hassan et al. 2022) was used to search the motor hotspot and estimate the resting motor threshold (RMT) using an automated closed-loop staircasing procedure. Target stimulation intensity was 130 %RMT (pre-sleep) or 100 %MSO (whichever was lower), resulting in average stimulation intensities of 128 %RMT (SD: 3 %RMT) or 95 %MSO (SD: 5%MSO), respectively. These comparably high intensities were chosen to ensure sufficiently large MEPs during the main experiment despite the known general suppression of MEPs during sleep (Avesani et al. 2008; Bergmann et al. 2012; Grosse et al. 2002; Salih et al. 2005) as well as the reduced coil output due to the 2m cable extension by which the coil was connected through a shielded cable guide to the stimulator outside the cabin. To avoid TMS discharge sound (“TMS click”) related sleep arousals, subjects wore in-ear headphones that delivered a continuous TMS-masking noise generated by the TAAC toolbox (Russo et al. 2022).

The inter-trial interval (ITI) between consecutive single TMS pulses was designed to range from ∼3 to ∼7 seconds, independent of sleep spindle occurrence (see Results for details). Before the start of the experiment, the BEST toolbox generated a trial vector with pseudorandomized experimental conditions, which was then “opportunistically” updated in real-time based on the number of ongoing spindle events in each set of six experimental conditions. The opportunistic strategy in the BEST toolbox was designed to operate in a way that if a planned condition was unavailable, but any of the next 5 planned conditions became possible, the BEST toolbox would trigger for that next possible condition and replace it in the sequence with the currently planned one. This ensured efficient use of available spindle events. In cases where none of the 6 consecutively planned conditions in a block occurred and the maximum ITI of 7 s was reached, the BEST toolbox would trigger a dummy pulse regardless of the planned condition. This approach allowed for flexible adaptation to the unpredictable nature of spindle events while maintaining a quasi-continuous sequence of stimulations within the specified ITI range that would prevent extreme ITI durations from influencing and confounding MEP amplitudes (Hassanzahraee et al., 2019).

### 2.5 Real-time EEG-TMS

The Real-Time Spindle Detector (RTSD) was used to perform real-time EEG analysis to detect sleep spindles and their instantaneous oscillatory phases, as previously described in detail (Hassan et al. 2022). In summary, EEG data was streamed in real-time to the bossdevice (sync2brain, Germany) that was controlled by the BEST toolbox (Hassan et al. 2022). To detect centroparietal spindles and their instantaneous oscillatory phases, a two-channel based spatial filter (bipolar montage) from EEG electrodes C3 and M2 (right mastoid) was used to extract the signal from the left sensorimotor cortex. EEG recorded during the adaptation nap from each subject was analyzed using YASA (Vallat and Walker 2021), i.e., an offline spindle detection software, to obtain the individual spindle band frequency and spindle frequency band (sigma) root mean square (RMS) power threshold required by RTSD for the experimental session, during which instantaneous oscillatory phase was estimated in real-time using the previously published *phastimate* algorithm (Zrenner et al. 2020). The RTSD (Hassan et al. 2022) operates by analyzing 520 ms segments of EEG montage every 10 ms, with a 98% overlap between consecutive windows. It employs two-pass FIR filters to create a broadband signal (EEG_bb_) using a 1-30 Hz passband, and a sigma band signal (EEG_σ_) using a passband centered on the individual’s spindle peak frequency ± 2 Hz. Four high-level signals are then derived: EEGσ RMS power, EEG_σ_ relative power, EEG_σ_ x EEG_bb_ correlation and EEG_bb_ instantaneous frequency. Spindles are detected when at least three of four derived signals exceed individualized thresholds (based on prior baseline sleep data), while also meeting specific duration criteria.

Real-time detection of SOs was merely done to target the spindle- and SO-free baseline NREM epochs conditions and was also performed in the bossdevice. The spatially filtered (i.e. re-referenced) C3-M2 EEG signal was bandpass filtered using two-pass, zero-phase, non-causal finite impulse response (FIR) filter (0.3-1.5 Hz bandpass; Hamming window passband ripple: 0.0194 with 53 dB stopband attenuation; Lower transition bandwidth: 0.20 Hz at -6 dB; Upper transition bandwidth: 1.6 Hz at -6 dB; Filter order: 500). SOs were detected when the bandpass signal exceeded the threshold of mean ± 2 times its standard deviation and the time interval between two subsequent zero crossings in opposite directions was between 0.1 and 1.5 seconds. While this approach diverges from standard offline methods that typically define SO duration as the full period between positive-to-negative zero crossings, it enabled earlier detection within each potential SO cycle, crucial for precisely timing our designed experimental conditions. YASA was used to extract the individual SO amplitude threshold from the adaptation nap data. Note that this comparably crude SO detection merely served to determine SO-free intervals in real-time and not to perform a detailed SO detection and characterization.

During the experimental sessions, PSG was monitored online to visually score the sleep stages using standard guidelines of the American Academy of Sleep Medicine (AASM) to manually start the real-time EEG triggered TMS at the start of NREM sleep stage 2. TMS intensity was gradually increased to ensure that the subject was not awakened by the sudden delivery of pulses. The total number of ramping pulses varied across subjects, ranging from 12 to 40 trials. All trials collected during the ramping process were discarded later.

### 2.6 Offline EEG analysis

Standard guidelines of AASM were used to visually score sleep stages using electrode C3 (0.5-40 Hz bandpass filtered) after excluding periods with TMS pulse artifacts and TMS-evoked K-complexes. Unless stated otherwise, all offline analyses were performed using the MNE-Python package (Gramfort et al. 2014) and FieldTrip toolbox (Oostenveld et al. 2011). EEG analyses were performed post-hoc to distinguish spindles co-occurring with SO from isolated spindles, to validate the performance of real-time EEG analyses, and to calculate the morphological parameters of detected spindles for trial-by-trial correlations with MEP amplitude. EEG data were segmented (−2.5 to -0.004 s relative to TMS), baseline corrected (−0.504 to −0.004 s, avoiding the TMS pulse artifact), and re-referenced to the common average of all EEG electrodes. A virtual channel representing the left sensorimotor cortex signal [C3 - M2 (right mastoid)] was added. EEG data prior to the TMS pulse (-1.5004 to -0.004 s) in the preprocessed EEG segments were further analyzed for offline detection of spindles co-occurring with SO using the slow-wave detection algorithm implemented in YASA (Vallat and Walker 2021) based on (Carrier et al. 2011; Massimini et al. 2004). All detected spindles that co-occurred with SOs were removed from further analysis as the trial count was insufficient (Mean: 37; SD: 8) to make a reliable comparison of isolated spindle conditions to those that co-occurred with SO.

The Pre-TMS EEG data were then further analyzed to verify that TMS was accurately delivered during the intended EEG-defined conditions. To demonstrate frequency-specificity to the targeted sleep spindles, time-frequency representations (TFR) were calculated for the pre-TMS period, for a period ranging from -1.5 to 1 s, with the post-TMS period replaced by zeros to prevent any TMS-related responses and artifacts from corrupting power estimates in the pre-TMS period. We applied the multitapers convolution method with a dynamic window length of 5 cycles of a given frequency, a step size of 20 ms, and a frequency resolution of 0.5 Hz, with a range of 1 to 35 Hz baseline corrected from [-0.5 to -0.4] s. To illustrate the topographical specificity of the targeted sleep spindle power, the topographical distribution of baseline corrected pre-TMS spindle power values was plotted per condition. Pre-TMS time-series were averaged across trials per subject and condition to demonstrate phase-specificity. Because the instantaneous phase of TMS delivery could not be calculated directly for the time point of stimulation due to signal corruption caused by TMS-evoked potentials and -related artifacts, it was estimated for each trial at one individual spindle cycle earlier (Bergmann et al. 2019). It is worth noting that the phase estimates do not reveal any hardware or software-related signal processing delays because they were compensated for prior to forecasting the instantaneous oscillatory phase in each trial.

The pre-TMS data (-1.004 to -0.004 s) from all the spindle trials were analyzed to calculate morphological parameters i.e., frequency, sigma power, instantaneous sigma amplitude at the time of TMS delivery, maximum sigma amplitude, spindle duration, and 1/f level (**Table S1**). The power spectrum obtained from the fast Fourier transform (FFT) of a 1 second pre-TMS data was used to calculate frequency and sigma power. The maximum and instantaneous sigma amplitude were obtained by calculating the sigma band power (previously described in RTSD; (Hassan et al. 2022). The duration of the spindle from its onset to the time of TMS delivery was calculated as the difference between time at TMS delivery and its nearest possible sample when the sigma RMS power was found to be less than the “entry RMS threshold” of mean + 1.15 SDs (standard deviation) of the baseline RMS power, i.e., adaptation nap sigma RMS power. The 1/f levels, i.e., average power of the aperiodic signal across the sigma frequency band, were determined by applying the “fitting oscillations & one-over f (FOOOF)” tool (Donoghue et al. 2020).

Preprocessed EEG data segments were averaged per condition (spindle-peak, spindle-falling flank, spindle-trough, spindle-rising flank, spindle-refractory period, no-spindle/SO/K-complex) time-locked to the TMS pulse. Subject-wise averages were then aggregated to obtain grand averages for each condition.

### 2.7 Offline EMG analysis

EMG data were analyzed using BEST and FieldTrip toolboxes (Hassan et al. 2022; Oostenveld et al. 2011). In summary, raw EMG data was epoch from [-50 to +100] ms around the TMS pulse. To rectify the DC offset, the data were demeaned relative to a pre-TMS data window of [-50 -5] ms. The amplitude of the MEP was computed by measuring the peak-to-peak value within a constant window of [15 50] ms after TMS pulse. MEP peak-to-peak amplitudes were additionally normalized block-wise as percent change from a running window average of 120 trials (across all conditions) and then averaged across blocks to take slow drifts in corticospinal excitability into account (**Figure S2**) (Bergmann et al. 2019; Thies et al. 2018; Zrenner et al. 2018).

### 2.8 Statistics

The independent variable was the targeted EEG-defined NREM sleep state, which was realized as a within-subject factor with the following six levels: spindle trough, spindle rising flank, spindle peak, spindle falling flank, spindle refractory period, and periods with no spindle, SO, or K-complex. The main dependent variable was corticospinal excitability as indexed by MEP amplitude. One-way rmANOVAs were conducted with post-hoc paired t-test where applicable. Statistical analyses were conducted using MATLAB (functions RMAOV1 and ttest), and a p-value < 0.05 was considered significant. Effect sizes for ANOVA (F-values) and *t* tests (Cohen’s *d*av, based on the averaged SD) are provided (Lakens, 2013). The electrode-wise MEP-spindle single trial correlations in supplementary analysis (**Figure S1**) were tested for significance using cluster-based permutation tests with correction for multiple comparisons (Oostenveld et al. 2011). Correlational analyses were performed on each channel to test for a linear relationship between trial-by-trial variations in normalized MEP amplitudes and spindle morphological parameters at the time of stimulation. Correlation coefficients were calculated separately for all spindle phase-triggered conditions in every subject, Fischer z-transformed, and were then tested channel-wise on the group level using two-sided paired t-test.

## 3 Results

During the experimental nap, where TMS was administered, subjects slept for an average of 97.3 ± 21 minutes. Average spindle density during the experimental nap was 4.8 ± 0.4 spindles per minute NREM sleep, and subjects received a total of 796 ± 84 TMS pulses, including dummy trials. On average, 86 ± 15 trials were acquired per experimental condition (spindle-peak, spindle-falling flank, spindle-trough, spindle-rising flank, spindle-refractory period, spindle-free), resulting in 74 ± 5 trials included in the analyses after exclusion of trials where spindles occurred concurrently with an SO. The median inter-trial interval between two consecutive TMS trials was 4.7 ± 1.2 s, with no statistical difference between conditions (paired t-test p > 0.7).

### 3.1 Sleep spindles phasically inhibit MEP corticospinal excitability

MEP amplitudes were modulated by sleep spindle phases (F_(5,95)_ =4.250, p=0.0016; **Figure 1C**). MEPs were smaller during the sleep spindle falling flank than during any other condition (falling vs trough: t_22_= 3.92, p = 0.00046, Cohen’s *d*av = 0.88; falling vs rising: t_22_= 3.34, p = 0.00344, Cohen’s *d*av = 0.71; falling vs peak: t_22_= 3.41, p = 0.00294, Cohen’s *d*av = 0.76; falling vs refractory: t_22_= 3.15, p = 0.002, Cohen’s *d*av = 0.67; falling vs spindle-free: t_22_= 3.59, p = 0.0009, Cohen’s *d*av = 0.77). All other comparisons were non-significant (p > 0.3). This means that corticospinal excitability was rhythmically suppressed during the falling flanks of the sleep spindle oscillation (by 12% on average) while remaining at levels comparable to the spindle-free baseline NREM periods during the spindle troughs, rising flanks, peaks, and refractory post-spindle periods. Sleep spindle phases also modulated normalized MEP amplitudes (**Figure S2**).

When ignoring spindle phase and averaging across all spindle conditions, MEP amplitude was still smaller than during the absence of spindle/SO/K-complex (spindle-free condition) in NREM sleep (t_22_= 1.96, p = 0.032, Cohen’s *d*av = 0.42; **Figure 1D**) and spindle refractory period (t_22_= 1.81, p = 0.043, Cohen’s *d*av = 0.39; **Figure 1D**). A comparison of the MEPs measured during the spindle refractory period and the baseline spindle-free oscillatory state in NREM sleep revealed no statistically significant difference (p > 0.6). By comparing the pre-sleep wakefulness cortical excitability with NREM spindle-free periods we found the MEP amplitude to be generally decreased (by 39% on average) during NREM sleep (t_22_= 3.74, p = 0.0007, Cohen’s *d*av = 0.80; **Figure 1E**).

### 3.2 Real-time EEG-triggered TMS successfully targeted sleep spindle and phases

The additional offline analyses of pre-TMS EEG confirmed that TMS was delivered at the intended EEG-defined brain-states and that no systematic confounds occurred (**Figure 2**) as evident by time-frequency responses (**Figure 2A**), topographical distributions of sleep spindle power (**Figure 2B**), estimated spindle phases at the time of actual TMS pulse delivery (**Figure 2C**), and time-locked averaged EEG signals (**Figure 2D**). According to these analyses, sleep spindles, spindle phases, and desynchronized oscillatory states were consistently targeted across subjects (mean circular standard deviation of 51° for all sleep spindle phase targeted conditions; ∼10 ms for an averaged 13 Hz sleep spindle), and adjacent frequencies or oscillatory activity from other sources, e.g., SO or broad frequency band noise did not confound the results.

**Figure 2:**
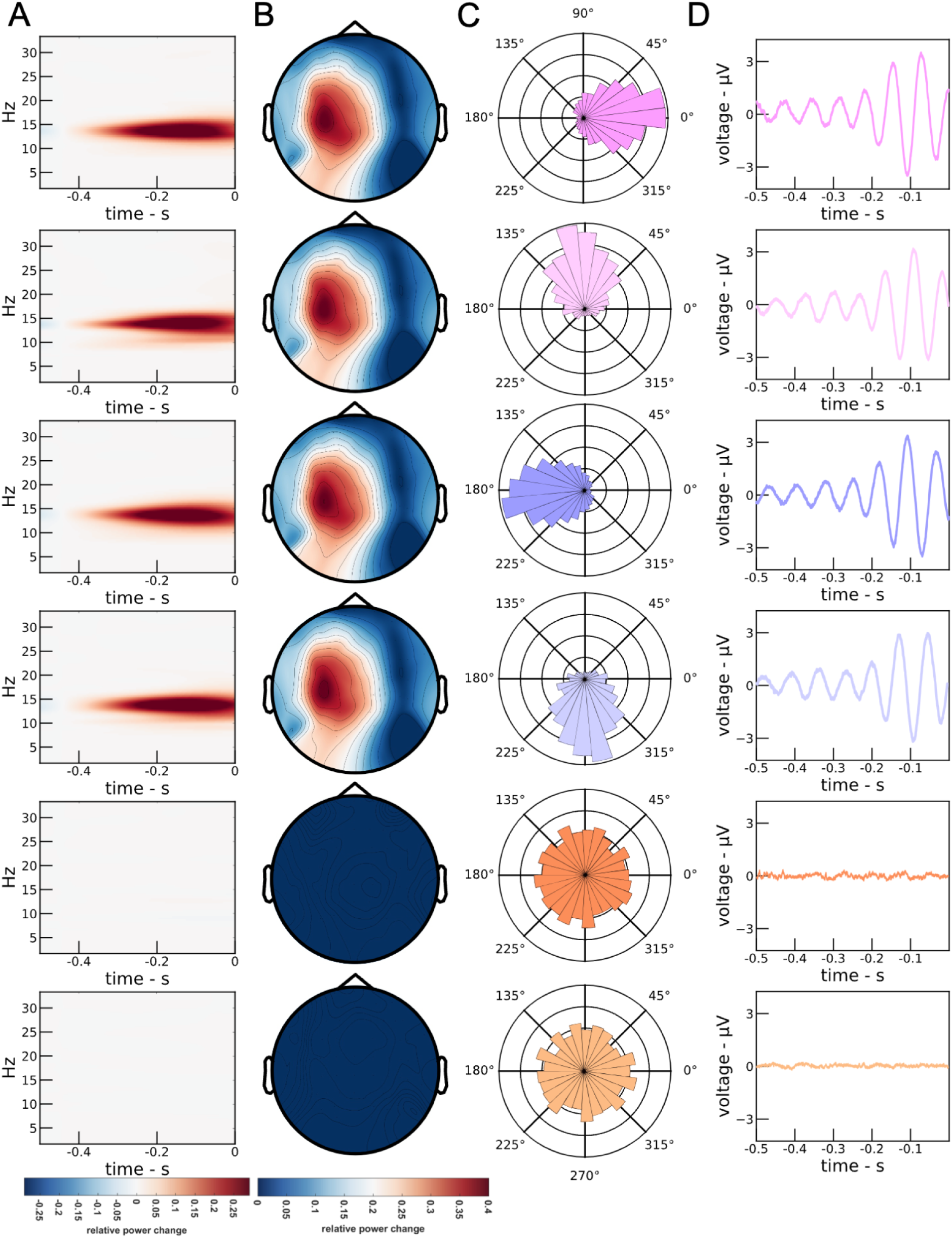
Sleep spindles and instantaneous phases were successfully targeted. **(A)** TFRs of oscillatory power in the C3-right mastoid signal prior to TMS (delivered at time point 0 ms). **(B)** Topographical maps of pre-TMS spindle power. **(C)** Phase histogram illustrating instantaneous phases in the C3-right mastoid signal one spindle cycle before TMS delivery. **(D)** Averaged time-locked C3-right mastoid signal relative to delivery of TMS (at 0 ms) highlighting sigma (12-15 Hz) activity in spindle conditions.

## Discussion

We report first empirical evidence that the sleep spindle reflects asymmetrical pulsed inhibition of corticospinal excitability. MEP amplitudes were inhibited during the falling flank of the spindle oscillation, relative to the desynchronized baseline NREM sleep state, when spindles/SOs/K-complexes were absent (spindle-free), spindle refractory periods, as well as spindle troughs, rising flanks, and peaks. Our findings suggests that sleep spindles do not simply exert a tonic inhibition of corticospinal excitability compared to spindle-free or spindle refractory periods, but a rhythmic suppression at a very limited part of the spindle oscillatory cycle, with no effect during the remaining phases. This study further confirms previous reports that corticospinal excitability is markedly suppressed during NREM sleep, as evidenced by lower MEP amplitudes compared to pre-sleep wakefulness. Unlike established for SOs in a previous study (Bergmann et al. 2012), there was no evidence linking corticospinal excitability (MEP amplitude) to spindle instantaneous amplitude nor any other morphological characteristic such as sigma oscillatory power, 1/f level, peak spindle frequency, spindle duration, or maximum instantaneous spindle amplitude.

### 3.3 Sleep spindle phase-dependent rhythmic inhibition of corticospinal excitability

Sleep spindle falling flanks, but not any of the other tested phase angles (troughs, rising flanks, and peaks), were associated with inhibition of corticospinal excitability relative to the spindle-free baseline NREM state, while no relative facilitation was observed during any of the experimental conditions. To date, no comparable data exists that provide insights into the phase-resolved excitability profile of sleep spindles, neither from human nor animal work. Therefore, we can only compare our findings to those of previous studies on other neuronal oscillations and their phase-dependent modulation of M1 corticospinal excitability. During NREM sleep, the SO up-state was shown to be associated with larger excitability than the SO down-state (Bergmann et al. 2012). Unfortunately, no MEPs were recorded during SO-free NREM sleep as baseline, so that a potential asymmetry and directionality of the SO phase-dependent modulatory effect could not be investigated. Also, rising and falling flanks of the SO were not probed, so that the actual excitability minima and maxima of the SO remain yet to be determined. During wakefulness, corticospinal excitability was found to be larger during the trough than peak of the sensorimotor mu-alpha (∼8-14 Hz) rhythm (Bergmann et al. 2019; Schaworonkow et al. 2019; Stefanou et al. 2018; Zrenner et al. 2018), later also qualified as asymmetric pulsed facilitation relative to a desynchronized mu-alpha-free background state (Bergmann et al. 2019). A recent study replicated mu-alpha phase-dependent effects on M1 cortical excitability (Wischnewski et al. 2022). Furthermore, also studies using post-hoc sorting of trials according to mu-alpha phase revealed a similar phasic modulation of corticospinal excitability (Ozdemir et al. 2022; Zrenner et al. 2022), while further refining the excitability maximum from trough to early rising phase of the mu-alpha cycle (Zrenner et al. 2023). Similarly, future studies may use either spindle-triggered random phase MEP measurements or phase-triggered MEPs at higher phase resolution to further narrow down the specific excitability minima/maxima for spindles. In any case, our findings appear to be in good agreement with the notion of an oscillatory phase-dependent modulation of corticospinal excitability in the human M1.

A direct comparison of the phasic excitability profiles of spindle and mu-alpha oscillations, which cover roughly the same frequency space during NREM sleep and wakefulness, respectively, and both show a topographical maximum over central sensorimotor regions, reveals interesting differences and similarities. In fact, the relative phase-excitability relationship appears highly similar for the two oscillations, while the asymmetry of the phasic modulation appears to be of opposite direction, resembling pulsed facilitation for mu-alpha versus pulsed inhibition for spindles. Whether this is due to their different neurophysiological mechanisms of generation or different excitability levels of the respective baseline periods during wakefulness and sleep, remains to be determined. Notably, both spindles and mu-alpha show a very different phase-excitability relationship than SOs, for which the excitability was larger during the surface EEG peak than trough, here reflecting the depolarized up-state and hyperpolarized down-state, respectively.

### 3.4 Spindle refractoriness has no effect on corticospinal excitability

Spindles are less likely to occur up to 3 s after the occurrence of the previous spindle (Antony et al. 2018), indicating a refractory period during which the neurophysiological mechanisms necessary for spindle generation may be dormant (Bal and McCormick 1996; Koupparis et al. 2013; Lüthi and McCormick 1998). Prior research even suggested that spindle refractoriness may be crucial for memory reactivation (Antony et al. 2018), as targeted memory reactivation had stronger effects on memory retention if applied after as compared to during the spindle refractory period (Antony et al. 2018). However, this refractory period does not seem to be related to a transient reduction in cortical excitability but rather excitability changes in thalamic neurons, as we did not observe a difference in MEP amplitudes between random periods of desynchronized baseline NREM spindle-free periods and periods of spindle refractoriness.

### 3.5 Sleep-dependent changes in corticospinal excitability

Our findings also corroborate the notion of a general NREM sleep-dependent suppression of corticospinal excitability, as reported by previous TMS studies during sleep. MEP amplitudes were generally found to be decreased during NREM sleep compared to pre-sleep wakefulness (Avesani et al. 2008; Bergmann et al. 2012; Grosse et al. 2002; Salih et al. 2005). This general transition from pre-sleep wakefulness to deep sleep has been linked to suppression of excitability at the spinal level, possibly also related to the characteristic neuromuscular blockade during NREM - and in particular REM sleep (Grosse et al. 2002) - but increased paired-pulse short latency intracortical inhibition (SICI) suggests a cortical contribution as well (Avesani et al. 2008; Salih et al. 2005).

### 3.6 Relevance of spindle morphology

We found no correlation between sleep spindle instantaneous amplitude and MEP amplitude. This result is not in line with previous studies, as a significant trial-by-trial correlation between the actual oscillatory SO amplitude and the evoked MEP amplitude was observed during sleep (Bergmann et al. 2012), and event-wise variations in spindle amplitude also predicted spindle-related BOLD response amplitude during concurrent EEG-fMRI measurements (Bergmann et al. 2012). Additionally, real-time EEG power triggered TMS also demonstrated that corticospinal excitability increased with the power of the sensorimotor mu-alpha rhythm (Thies et al. 2018). Further analysis also revealed no relationship between MEP amplitude and any of the other calculated spindle morphological parameters including sigma power, frequency, duration, 1/f level, or maximum spindle amplitude. We assumed that MEP amplitudes would, at least during the spindle falling-flank phase angle experimental condition, correlate with the trial-by-trial variation in spindle instantaneous amplitude or other morphological parameters because MEPs in this condition were significantly suppressed compared to all other conditions. It is possible that the amplitude threshold used for spindle detection did reduce their variance to a degree that the remaining spindle amplitude differences were too small to impact MEP amplitude.

### 3.7 Potential mechanisms mediating sleep spindle phase-dependent inhibition of corticospinal excitability

Sleep spindles are generated in thalamic nuclei as an interplay between GABA-ergic neurons in the thalamic reticular nucleus (TRN) and glutamatergic thalamocortical (TC) relay neurons, which project the spindle patterns into the neocortex, while both TRN and TC cells receive excitatory feedback from corticothalamic (CT) cells in return, allowing the neocortex to influence spindle generation, in particular during their initiation and termination (Fernandez and Lüthi 2020; Gardner et al. 2013). Accordingly, research in rodents using calcium imaging and LFP recordings has linked spindle generation to the activity of parvalbumin (PV+)-positive GABAergic inhibitory interneurons in the TRN and other thalamic nuclei (Fernandez et al. 2018; Raven and Aton 2021; Thankachan et al. 2019). But also in the neocortex, a specific activation pattern of different neuron types has been observed during spindles and SOs in NREM sleep (Klinzing et al. 2019). The average firing rate during spindles was found to be increased not only for excitatory glutamatergic Pyramidal (PYR) cells (Brécier et al. 2022; Niethard et al. 2018) but even much more so for PV+ inhibitory interneurons (Brécier et al. 2022; Niethard et al. 2018; Peyrache et al. 2011), while firing rates of somatostatin (SOM+)-positive GABAergic inhibitory interneurons decreased (Brécier et al. 2022; Niethard et al. 2018) and those of vasoactive intestinal polypeptide (VIP)-positive interneurons remained unchanged (Brécier et al. 2022). Notably, in contrast to the strong PV+ involvement during spindles, Niethard et al. (2018) found SOs to be associated in particular with the firing of SOM+ cells, while spindles nested in SO up-states showed strong PV+ activity, reduced SOM+ activity, and a 3-fold increase in PYR activity compared to the respective solitary events. These observations are corroborated by causal optogenetic and chemogenetic manipulations, which have shown that activation/deactivation of SOM+ cells increases/decreases slow wave activity (SWA), whereas activation of PV+ cells decreases SWA (Funk et al. 2017). This particular constellation of excitatory and inhibitory input to PYR cells has been suggested to facilitate processes of synaptic rescaling during systems memory consolidation, as the weak dendritic inhibition by SOM+ cells enables strong postsynaptic activation of PYR cells, while their strong perisomatic inhibition by PV+ cells prevents them from firing (Klinzing et al. 2019; Niethard et al. 2018). Unfortunately, the employed calcium imaging techniques do not provide the temporal resolution to provide spindle phase-resolved profiles of PYR, PV+, SOM+, and VIP+ activity, but electrophysiological recordings may provide that information in future studies. Single- and multi-unit recordings in humans have confirmed the increased spiking activity of both putative PYR and inhibitory interneuron (IN) cells (without the possibility for further distinction into PV+, SOM+, and VIP+) during spindles in the local field potential (LFP), in particular those nested in SO up-states (Dickey et al. 2021). These recordings also revealed phase locking of both neuron types to the spindle oscillation as well as an increased co-firing of neuron pairs (within STDP-relevant 25ms windows) of both types during the spindle, again particularly during spindle-SO events.

Increases of PV+ interneuron activity have also been implicated more generally in the generation and regulation of sleep-related neuronal oscillations (Raven and Aton 2021), including hippocampal ripples (Forro et al. 2015; Somogyi et al. 2014; Szabo et al. 2022) and gamma oscillations (Buzsáki and Wang 2012). Importantly, both cortical gamma and hippocampal ripples appear to be phase-modulated by spindle oscillations. Analyzing simultaneous EEG and Magnetoencephalography (MEG) results, Ayoub et al. (2012) found cortical gamma power (MEG) to be more pronounced during surface-positive spindles (EEG), whereas hippocampal ripples were found to be phase-locked to spindle peaks in stereoelectroencephalography (sEEG) recordings in partial seizure patients (Jiang et al. 2019) and to spindle troughs in intracranial-EEG (iEEG) recordings in epilepsy patients (Staresina et al. 2015) as well as during local fields potential (LFP) recordings in rodents (Sirota et al. 2003). While the different preferred phase angles of these phase-amplitude coupling (PAC) results may be difficult to reconcile at first glance, one must keep in mind that they are not directly comparable due to differences in electrophysiological recording modalities as well as the position of recording and reference electrodes. These results, however, converge on the notion that spindles phasically modulate beta and faster oscillations. And while PAC itself is agnostic to whether it is mediated by rhythmic suppression or facilitation, the increased PV+ interneuron activity together with the phase-specific suppression of corticospinal excitability revealed in the current paper, suggests spindle phase-related rhythmic inhibition rather than rhythmic facilitation of neuronal activity. Future studies may use paired-pulse TMS protocols, such as GABA-A-receptor mediated SICI, to investigate the inhibitory mechanism underlying the observed phasic modulation of MEPs, as done for the sensorimotor mu-alpha rhythm (Bergmann et al. 2019).

### 3.8 A route towards spindle phase-dependent plasticity and probing their causal role in sleep-dependent memory consolidation

Beside advancing our fundamental understanding of spindle physiology, the unique experimental setup we established and the resulting characterization of a clear spindle phase-excitability relationship have also direct implications for the development of novel EEG-triggered TMS protocols to investigate both spindle phase-dependent plasticity and the causal role of spindles in memory reactivation and consolidation. To tap into the mechanism of phase-dependent plasticity (Bergmann and Born 2018), TMS burst need to be delivered repetitively to the most (vs. least) excitable phase of an oscillation, as repeatedly demonstrated for the sensorimotor mu-alpha rhythm during wake, where repeated delivery of 100-200 Hz TMS bursts to M1 during the more excitable trough of the oscillation produced a lasting long-term potentiation (LTP)-like increase in corticospinal excitability, whereas delivery of the same burst to the less excitable mu-alpha peaks did not, and random phase delivery of 200 Hz burst even caused long-term depression (LTP)-like decreases (Baur et al. 2020; Baur et al. 2022; Zrenner et al. 2018). Based on the observed phase-excitability relationship of spindle oscillations, one may therefore speculate that the repeated application of gamma-/ripple-like TMS bursts during spindles targeting the phase of maximum inhibition (for our EEG montage corresponding to the falling flanks of the oscillation) would cause LTD-like MEP decreases, while targeting the troughs may cause LTP-like increases. The latter assumption would be based on the notion that spindle troughs descriptively showed the largest MEPs (even though not different from baseline or any of the other conditions despite falling flanks) but more importantly that this phase immediately follows the strong suppression during the falling flank and thereby potentially represents a state of release from inhibition (or dis-inhibition) that may facilitate LTP (Cash et al. 2016).

It can be suggested that sleep spindle trough (least inhibitory) triggered rTMS may lead to LTP-like effects similar to the previously reported mu-alpha trough triggered (most excitable) rTMS of M1 that lead to LTP-like effects in wakefulness (Zrenner et al. 2018). These mu-alpha dependent LTP-like effects were attributed to the disinhibition potentially associated with the facilitatory effects of motor cortical neurons at the trough following the least facilitatory peaks, given the general relevance of disinhibition for TMS-related LTP-like plasticity (Cash et al. 2016). Similar to the recently reported mu-alpha random phase triggered 200 Hz bursts of rTMS over M1 that induce LTD-like effects during wakefulness (Baur et al. 2022), tonically (i.e., random phase) triggered rTMS during thalamocortical spindle may also result in LTD-like effects on corticospinal excitability. Moreover, repeatedly targeting spindles or even specific spindle phases with TMS may allow researchers to deliberately interfere with processes of memory reactivation and associated synaptic rescaling in the neocortex as part of the hippocampo-neocortical dialogue mediated by SO-Spindle-ripple PAC (Diekelmann and Born 2010; Rasch and Born 2013; Staresina et al. 2015) and thereby demonstrate the causal role of sleep spindles in systems memory consolidation.

### 3.9 Limitations

This study is the first one of its kind, using real-time EEG spindle phase-triggered targeting of M1 with TMS during human NREM sleep to provide a phase-resolved characterization of the corticospinal excitability pattern associated with sleep spindles. This demanding methodological setup naturally comes with a number of limitations. Firstly, unlike established offline spindle detection algorithms that can use information of entire spindle waveforms and even entire sleep recordings, the dedicated real-time spindle detection (RTSD) algorithm (Hassan et al. 2022) we used is causal by design to work in real-time, meaning that it can only use the data recorded up to the current point in time. Therefore, spindles can only be detected after they have already emerged to a sufficient extent and meet certain amplitude criteria. They are thus detected and targeted more often in their second half (the waning phase of their amplitude), while their first half (with waxing amplitude) remains largely unstudied. Second, we sampled MEPs from only four spindle phase angles (trough, rising flank, peak, and falling flank), and it is well possible that additional or even stronger modulation occurs during phase angles not sampled in this study. Future studies using spindle event-triggered but phase-random targeting with post hoc phase-sorting may reveal a more fine-grained resolution of the phasic excitability dynamics, as in (Zrenner et al. 2023). Third, spindles and SOs frequently co-occur in deep NREM sleep (Staresina et al. 2015; Mölle et al. 2002; Mölle et al. 2011), and the precise spindle-SO phase-amplitude coupling has been shown to affect neuronal (co-)firing (Dickey et al. 2021) and the specific pattern of excitatory and inhibitory neuronal involvement (Niethard et al. 2018). This is thought to play an important role in memory consolidation (Hahn et al. 2020; Helfrich et al. 2018; Latchoumane et al. 2017), likely via the mechanism of phase-dependent plasticity (Bergmann and Born 2018). Comparing phasic corticospinal excitability modulations during isolated spindles with those nested in SO up-states would thus have been highly interesting but was unfortunately not possible due to very little deep NREM sleep in the current study and thus an insufficient number of spindles co-occurring with SOs. Fourth, the observed modulation of corticospinal excitability by spindle phase may be specific to M1 and not translate to other cortical regions and other outcome measures or do so but with shifts in the specific phase angles showing maximal effects, due to the traveling wave like properties of spindles and SOs and the resulting phase differences between cortical regions (Dickey et al. 2021; Massimini et al. 2004).

## 4 Conclusion

We provide the first in-human demonstration of the feasibility of real-time sleep spindle phase-triggered TMS and of a phasic modulation of corticospinal excitability by the sleep spindle. Our findings are best explained by sleep spindles exerting an asymmetrical “pulsed inhibition” on corticospinal excitability, complementing previous work demonstrating a phasic modulation of corticospinal excitability by the sleep SO (Bergmann et al. 2012) as well as asymmetrical “pulsed facilitation” by the sensorimotor mu-alpha rhythm during wakefulness (Bergmann et al. 2019; Zrenner et al. 2018). Utilizing this technique, future work should further characterize the excitation-inhibition profile of sleep spindles, investigate their interaction with SOs, and test their role as windows of phase-dependent synaptic plasticity as well as their causal relevance for memory reactivation in the context of systems memory consolidation.

## Acknowledgments

T.O.B. received support from funding from the Boehringer Ingelheim Foundation (BIF) and the German Research Foundation (DFG Grant 362546008); C.Z. received support from the German Federal Ministry for Economic Affairs and Energy through an EXIST Transfer of Research (grant no. 03EFJBW169); U.Z. received support from the European Research Council (ERC Synergy) under the European Union’s Horizon 2020 research and innovation program (ConnectToBrain; grant agreement No. 810377) and from the German Research Foundation (DFG Grant 468645090).

## Code and data availability

The code and data that support the findings of this study are available from the main author, U.H., upon request.

## Author contributions

U.H., T.O.B., and U.Z. conceived the study and supervised the project. U.H. and T.O.B. planned the experiments. U.H., P.O., and M.A.M., conducted the experiments. U.H. performed analysis, prepared figures, and wrote the main draft of the manuscript. U.H., T.O.B., and U.Z. reviewed and finalized the manuscript. All authors critically revised, read and approved the manuscript.

## Supplementary Material

### No effect of spindle morphology on MEP corticospinal excitability

Trial-by-trial variations in spindle instantaneous amplitude at the time of TMS, sigma oscillatory power, 1/f level, spindle peak frequency, duration of spindle at the time of TMS, and maximum instantaneous spindle amplitude did not correlate with trial-by-trial variations in MEP amplitude. The correlations were extremely weak, statistically insignificant (p > 0.05) with r values ranging from -0.1 to 0.1, and topographies revealed that they did not emerge from similar cortical areas.

**Figure S1:**
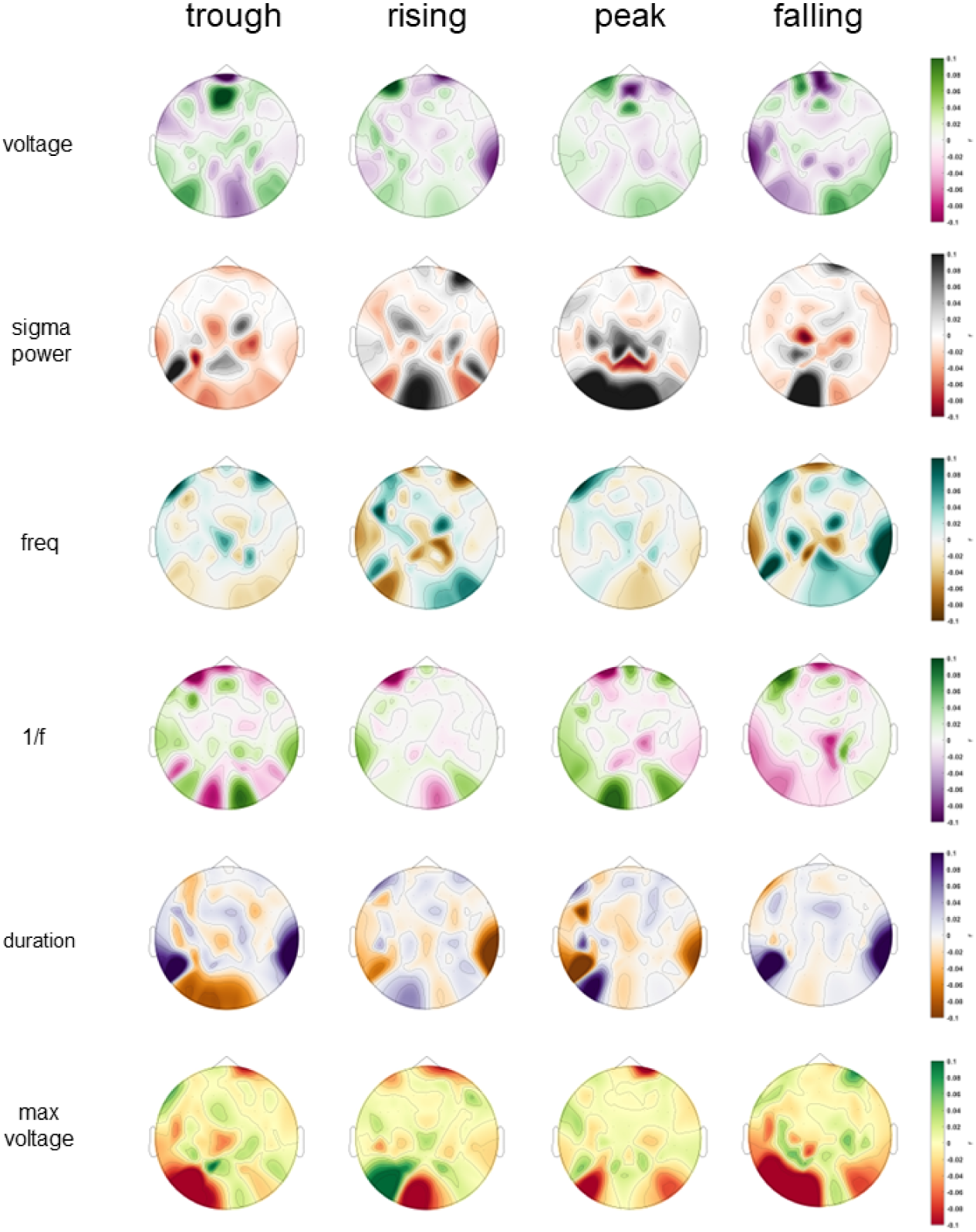
Correlational topographic maps of trial-by-trial variation in MEP amplitude and spindle morphological characteristics. Topographies reveal no significant difference of correlation (**r**) (p > 0.05) between the trial-by-trial variations of MEP amplitude and spindle morphological properties (amplitude/voltage, sigma power, frequency, 1/f level, duration, and maximum amplitude/voltage from onset).

**Figure S2:**
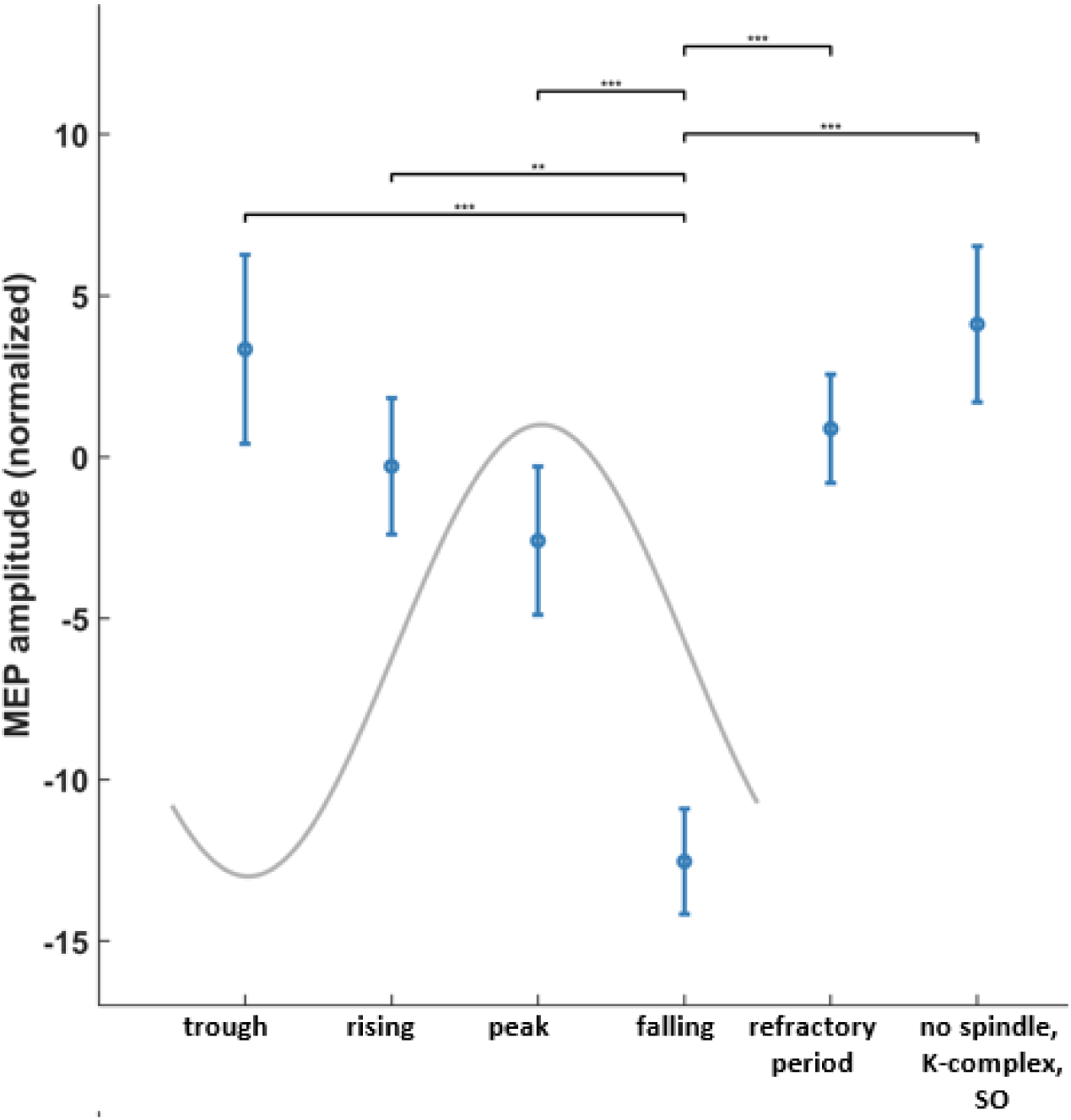
Thalamocortical spindle phase modulates normalized MEPs revealing a rhythmic inhibition of motor corticospinal excitability at the falling flank (see methods; offline EMG analysis).

**Supplementary Table S1:** Morphological properties of detected spindles (M ± SD).

| Subject No. | Voltage (uV) | Maximum Voltage (uV) | Frequency (Hz) | 1/f exponent-z | Duration (seconds) | Sigma power-z |
| --- | --- | --- | --- | --- | --- | --- |
| 1 | 3.1±0.5 | 4.7±0.9 | 13.4±1.8 | 0.56±0.13 | 0.64±0.08 | 0.74±0.06 |
| 2 | 2.9±0.3 | 4.3±0.7 | 14.6±2.2 | 0.41±0.08 | 0.76±0.09 | 0.89±0.07 |
| 3 | 2.4±0.8 | 4.5±1.2 | 12.2±1.4 | 0.65±0.14 | 0.48±0.06 | 0.56±0.02 |
| 4 | 1.5±0.9 | 3.7±1.3 | 14.1±2.0 | 0.48±0.06 | 0.69±0.08 | 0.81±0.05 |
| 5 | 2.8±0.5 | 5.0±0.9 | 13.8±1.9 | 0.62±0.15 | 0.55±0.07 | 0.64±0.04 |
| 6 | 3.5±0.7 | 5.8±1.1 | 11.8±1.3 | 0.35±0.06 | 0.73±0.09 | 0.92±0.08 |
| 7 | 4.3±1.1 | 6.2±1.5 | 14.5±2.2 | 0.54±0.08 | 0.41±0.05 | 0.51±0.03 |
| 8 | 2.9±0.7 | 4.8±1.1 | 12.9±1.6 | 0.68±0.13 | 0.66±0.07 | 0.78±0.07 |
| 9 | 3.3±1.0 | 5.1±1.4 | 13.1±1.7 | 0.47±0.10 | 0.78±0.09 | 0.66±0.02 |
| 10 | 1.9±0.5 | 3.5±0.9 | 14.7±2.3 | 0.59±0.14 | 0.52±0.06 | 0.86±0.09 |
| 11 | 4.1±1.2 | 6.0±1.6 | 12.0±1.3 | 0.32±0.05 | 0.72±0.08 | 0.59±0.03 |
| 12 | 2.8±0.7 | 4.6±1.1 | 14.3±2.1 | 0.51±0.07 | 0.60±0.06 | 0.90±0.09 |
| 13 | 1.2±0.7 | 2.9±1.1 | 12.7±1.5 | 0.66±0.12 | 0.75±0.09 | 0.71±0.03 |
| 14 | 1.9±0.8 | 3.6±1.2 | 13.5±1.8 | 0.38±0.07 | 0.45±0.05 | 0.48±0.01 |
| 15 | 4.2±1.5 | 6.3±1.9 | 11.3±1.2 | 0.60±0.10 | 0.70±0.09 | 0.85±0.06 |
| 16 | 5.3±1.7 | 7.4±2.1 | 14.0±2.0 | 0.44±0.09 | 0.57±0.06 | 0.61±0.01 |
| 17 | 3.4±0.7 | 5.2±1.1 | 13.2±1.7 | 0.57±0.09 | 0.67±0.09 | 0.76±0.04 |
| 18 | 3.9±0.9 | 5.7±1.3 | 12.5±1.5 | 0.53±0.12 | 0.61±0.08 | 0.53±0.02 |
| 19 | 3.2±0.8 | 5.0±1.2 | 14.2±2.1 | 0.50±0.11 | 0.63±0.07 | 0.82±0.08 |
| 20 | 3.4±0.6 | 5.1±1.0 | 13.7±1.9 | 0.63±0.11 | 0.58±0.07 | 0.69±0.05 |
| Mean±SD | 3.1±0.8 | 4.9±1.2 | 13.3±1.7 | 0.5±0.1 | 0.6±0.07 | 0.7±0.04 |

## References

Antony, J. W., Piloto, L., Wang, M., Pacheco, P., Norman, K. A., & Paller, K. A. (6 2018). Sleep Spindle Refractoriness Segregates Periods of Memory Reactivation. Current Biology: CB, 28, 1736–1743.e4.

Avesani, M., Formaggio, E., Fuggetta, G., Fiaschi, A., & Manganotti, P. (5 2008). Corticospinal excitability in human subjects during nonrapid eye movement sleep: single and paired-pulse transcranial magnetic stimulation study. Experimental Brain Research. Experimentelle Hirnforschung. Experimentation Cerebrale, 187, 17–23.

Ayoub, A., Mölle, M., Preissl, H., & Born, J. (2012). Grouping of MEG gamma oscillations by EEG sleep spindles. NeuroImage, 59(2), 1491–1500.

Bal, T., & McCormick, D. A. (8 1996). What Stops Synchronized Thalamocortical Oscillations? Neuron, 17, 297–308.

Barthó, P., Slézia, A., Mátyás, F., Faradzs-Zade, L., Ulbert, I., Harris, K. D., & Acsády, L. (6 2014). Ongoing Network State Controls the Length of Sleep Spindles via Inhibitory Activity. In Neuron (Vol. 82, pp. 1367–1379). 10.1016/j.neuron.2014.04.046

Baur, D., Ermolova, M., Souza, V. H., Zrenner, C., & Ziemann, U. (11 2022). Phase-amplitude coupling in high-gamma frequency range induces LTP-like plasticity in human motor cortex: EEG-TMS evidence. Brain Stimulation. 10.1016/j.brs.2022.11.003

Baur, D., Galevska, D., Hussain, S., Cohen, L. G., Ziemann, U., & Zrenner, C. (11 2020). Induction of LTD-like corticospinal plasticity by low-frequency rTMS depends on pre-stimulus phase of sensorimotor μ-rhythm. Brain Stimulation, 13, 1580–1587.

Bergmann, T. O., & Born, J. (1 2018). Phase-Amplitude Coupling: A General Mechanism for Memory Processing and Synaptic Plasticity? Neuron, 97, 10–13.

Bergmann, T. O., Lieb, A., Zrenner, C., & Ziemann, U. (12 2019). Pulsed Facilitation of Corticospinal Excitability by the Sensorimotor μ-Alpha Rhythm. The Journal of Neuroscience: The Official Journal of the Society for Neuroscience, 39, 10034–10043.

Bergmann, T. O., Mölle, M., Diedrichs, J., Born, J., & Siebner, H. R. (2 2012). Sleep spindle-related reactivation of category-specific cortical regions after learning face-scene associations. NeuroImage, 59, 2733–2742.

Bergmann, T. O., Molle, M., Schmidt, M. A., Lindner, C., Marshall, L., Born, J., & Siebner, H. R. (1 2012). EEG-Guided Transcranial Magnetic Stimulation Reveals Rapid Shifts in Motor Cortical Excitability during the Human Sleep Slow Oscillation. Journal of Neuroscience, 32, 243–253.

Bonjean, M., Baker, T., Lemieux, M., Timofeev, I., Sejnowski, T., & Bazhenov, M. (6 2011). Corticothalamic Feedback Controls Sleep Spindle Duration In Vivo. Journal of Neuroscience, 31, 9124–9134.

Brécier, A., Borel, M., Urbain, N., & Gentet, L. J. (6 2022). Vigilance and Behavioral State-Dependent Modulation of Cortical Neuronal Activity throughout the Sleep/Wake Cycle. In The Journal of Neuroscience (Vol. 42, pp. 4852–4866). 10.1523/JNEUROSCI.1400-21.2022

Buzsáki, G., & Wang, X.-J. (2012). Mechanisms of Gamma Oscillations. Annual Review of Neuroscience, 35(Volume 35, 2012), 203–225.

Carrier, J., Viens, I., Poirier, G., Robillard, R., Lafortune, M., Vandewalle, G., Martin, N., Barakat, M., Paquet, J., & Filipini, D. (2 2011). Sleep slow wave changes during the middle years of life. The European Journal of Neuroscience, 33, 758–766.

Cash, R. F. H., Murakami, T., Chen, R., Thickbroom, G. W., & Ziemann, U. (1 2016). Augmenting Plasticity Induction in Human Motor Cortex by Disinhibition Stimulation. Cerebral Cortex, 26, 58–69.

Choi, J., & Jun, S. C. (3 2022). Spindle-targeted acoustic stimulation may stabilize an ongoing nap. Journal of Sleep Research. 10.1111/jsr.13583

Choi, J., Won, K., & Jun, S. C. (2019). Acoustic Stimulation Following Sleep Spindle Activity May Enhance Procedural Memory Consolidation During a Nap. IEEE Access, 7, 56297–56307.

Destexhe, A., Bal, T., McCormick, D. A., & Sejnowski, T. J. (9 1996). Ionic mechanisms underlying synchronized oscillations and propagating waves in a model of ferret thalamic slices. Journal of Neurophysiology, 76, 2049–2070.

Dickey, C. W., Sargsyan, A., Madsen, J. R., Eskandar, E. N., Cash, S. S., & Halgren, E. (12 2021). Travelling spindles create necessary conditions for spike-timing-dependent plasticity in humans. Nature Communications, 12, 1027.

Diekelmann, S., & Born, J. (2 2010). The memory function of sleep. Nature Reviews. Neuroscience, 11, 114–126.

Donoghue, T., Haller, M., Peterson, E. J., Varma, P., Sebastian, P., Gao, R., Noto, T., Lara, A. H., Wallis, J. D., Knight, R. T., Shestyuk, A., & Voytek, B. (12 2020). Parameterizing neural power spectra into periodic and aperiodic components. Nature Neuroscience, 23, 1655–1665.

Feldman, D. E. (8 2012). The Spike-Timing Dependence of Plasticity. Neuron, 75, 556–571.

Fernandez, L. M. J., & Lüthi, A. (4 2020). Sleep Spindles: Mechanisms and Functions. Physiological Reviews, 100, 805–868.

Fernandez, L. M. J., Vantomme, G., Osorio-Forero, A., Cardis, R., Béard, E., & Lüthi, A. (12 2018). Thalamic reticular control of local sleep in mouse sensory cortex. eLife, 7. 10.7554/eLife.39111

Forro, T., Valenti, O., Lasztoczi, B., & Klausberger, T. (5 2015). Temporal Organization of GABAergic Interneurons in the Intermediate CA1 Hippocampus During Network Oscillations. Cerebral Cortex, 25, 1228–1240.

Funk, C. M., Peelman, K., Bellesi, M., Marshall, W., Cirelli, C., & Tononi, G. (2017). Role of Somatostatin-Positive Cortical Interneurons in the Generation of Sleep Slow Waves. The Journal of Neuroscience: The Official Journal of the Society for Neuroscience, 37(38), 9132–9148.

Gardner, R. J., Hughes, S. W., & Jones, M. W. (11 2013). Differential Spike Timing and Phase Dynamics of Reticular Thalamic and Prefrontal Cortical Neuronal Populations during Sleep Spindles. Journal of Neuroscience, 33, 18469–18480.

Gramfort, A., Luessi, M., Larson, E., Engemann, D. A., Strohmeier, D., Brodbeck, C., Parkkonen, L., & Hämäläinen, M. S. (2 2014). MNE software for processing MEG and EEG data. NeuroImage, 86, 446–460.

Grosse, P., Khatami, R., Salih, F., Kuhn, A., & Meyer, B.-U. (12 2002). Corticospinal excitability in human sleep as assessed by transcranial magnetic stimulation. Neurology, 59, 1988–1991.

Hahn, M. A., Heib, D., Schabus, M., Hoedlmoser, K., & Helfrich, R. F. (6 2020). Slow oscillation-spindle coupling predicts enhanced memory formation from childhood to adolescence. eLife, 9. 10.7554/eLife.53730

Hassan, U., Feld, G. B., & Bergmann, T. O. (9 2022). Automated real-time EEG sleep spindle detection for brain-state-dependent brain stimulation. Journal of Sleep Research. 10.1111/jsr.13733

Hassan, U., Pillen, S., Zrenner, C., & Bergmann, T. O. (1 2022). The Brain Electrophysiological recording & STimulation (BEST) toolbox. Brain Stimulation, 15, 109–115.

Hassanzahraee, M., Zoghi, M., & Jaberzadeh, S. (12 2019). Longer Transcranial Magnetic Stimulation Intertrial Interval Increases Size, Reduces Variability, and Improves the Reliability of Motor Evoked Potentials. Brain Connectivity, 9, 770–776.

Helfrich, R. F., Mander, B. A., Jagust, W. J., Knight, R. T., & Walker, M. P. (1 2018). Old Brains Come Uncoupled in Sleep: Slow Wave-Spindle Synchrony, Brain Atrophy, and Forgetting. Neuron, 97, 221–230.e4.

Jiang, X., Gonzalez-Martinez, J., & Halgren, E. (11 2019). Posterior Hippocampal Spindle Ripples Co-occur with Neocortical Theta Bursts and Downstates-Upstates, and Phase-Lock with Parietal Spindles during NREM Sleep in Humans. The Journal of Neuroscience: The Official Journal of the Society for Neuroscience, 39, 8949–8968.

Klinzing, J. G., Niethard, N., & Born, J. (10 2019). Mechanisms of systems memory consolidation during sleep. Nature Neuroscience, 22, 1598–1610.

Koupparis, A. M., Kokkinos, V., & Kostopoulos, G. K. (1 2013). Spindle Power Is Not Affected after Spontaneous K-Complexes during Human NREM Sleep. PloS One, 8, e54343.

Langdon, A. J., Breakspear, M., & Coombes, S. (12 2012). Phase-locked cluster oscillations in periodically forced integrate-and-fire-or-burst neuronal populations. Physical Review E, 86, 061903.

Latchoumane, C.-F. V., Ngo, H.-V. V., Born, J., & Shin, H.-S. (2017). Thalamic spindles promote memory formation during sleep through triple phase-locking of cortical, thalamic, and hippocampal rhythms. Neuron, 95(2), 424–435.e6.

Lustenberger, C., Boyle, M. R., Alagapan, S., Mellin, J. M., Vaughn, B. V., & Fröhlich, F. (8 2016). Feedback-Controlled Transcranial Alternating Current Stimulation Reveals a Functional Role of Sleep Spindles in Motor Memory Consolidation. Current Biology: CB, 26, 2127–2136.

Lüthi, A., & McCormick, D. A. (7 1999). Modulation of a pacemaker current through Ca2+-induced stimulation of cAMP production. In Nature Neuroscience (Vol. 2, pp. 634–641). 10.1038/10189

Lüthi, A., & McCormick, D. A. (3 1998). Periodicity of Thalamic Synchronized Oscillations: the Role of Ca2+-Mediated Upregulation of Ih. In Neuron (Vol. 20, pp. 553–563). 10.1016/S0896-6273(00)80994-0

Massimini, M., Huber, R., Ferrarelli, F., Hill, S., & Tononi, G. (8 2004). The Sleep Slow Oscillation as a Traveling Wave. Journal of Neuroscience, 24, 6862–6870.

Mölle, M., Bergmann, T. O., Marshall, L., & Born, J. (10 2011). Fast and Slow Spindles during the Sleep Slow Oscillation: Disparate Coalescence and Engagement in Memory Processing. In Sleep (Vol. 34, pp. 1411–1421). 10.5665/SLEEP.1290

Mölle, M., Marshall, L., Gais, S., & Born, J. (2002). Grouping of spindle activity during slow oscillations in human non-rapid eye movement sleep. The Journal of Neuroscience: The Official Journal of the Society for Neuroscience, 22(24), 10941–10947.

Niethard, N., Brodt, S., & Born, J. (5 2021). Cell-Type-Specific Dynamics of Calcium Activity in Cortical Circuits over the Course of Slow-Wave Sleep and Rapid Eye Movement Sleep. The Journal of Neuroscience: The Official Journal of the Society for Neuroscience, 41, 4212–4222.

Niethard, N., Burgalossi, A., & Born, J. (9 2017). Plasticity during Sleep Is Linked to Specific Regulation of Cortical Circuit Activity. Frontiers in Neural Circuits, 11. 10.3389/fncir.2017.00065

Niethard, N., Hasegawa, M., Itokazu, T., Oyanedel, C. N., Born, J., & Sato, T. R. (10 2016). Sleep-Stage-Specific Regulation of Cortical Excitation and Inhibition. Current Biology: CB, 26, 2739–2749.

Niethard, N., Ngo, H.-V. V., Ehrlich, I., & Born, J. (9 2018). Cortical circuit activity underlying sleep slow oscillations and spindles. Proceedings of the National Academy of Sciences, 115. 10.1073/pnas.1805517115

Oostenveld, R., Fries, P., Maris, E., & Schoffelen, J.-M. (2011). FieldTrip: Open source software for advanced analysis of MEG, EEG, and invasive electrophysiological data. Computational Intelligence and Neuroscience, 2011, 156869.

Ozdemir, R. A., Kirkman, S., Magnuson, J. R., Fried, P. J., Pascual-Leone, A., & Shafi, M. M. (12 2022). Phase matters when there is power: Phasic modulation of corticospinal excitability occurs at high amplitude sensorimotor mu-oscillations. Neuroimage: Reports, 2, 100132.

Peyrache, A., Battaglia, F. P., & Destexhe, A. (10 2011). Inhibition recruitment in prefrontal cortex during sleep spindles and gating of hippocampal inputs. Proceedings of the National Academy of Sciences, 108, 17207–17212.

Rasch, B., & Born, J. (4 2013). About Sleep’s Role in Memory. Physiological Reviews, 93, 681–766.

Raven, F., & Aton, S. J. (9 2021). The Engram’s Dark Horse: How Interneurons Regulate State-Dependent Memory Processing and Plasticity. Frontiers in Neural Circuits, 15. 10.3389/fncir.2021.750541

Rosanova, M. (10 2005). Pattern-Specific Associative Long-Term Potentiation Induced by a Sleep Spindle-Related Spike Train. Journal of Neuroscience, 25, 9398–9405.

Rossi, S., Hallett, M., Rossini, P. M., & Pascual-Leone, A. (8 2011). Screening questionnaire before TMS: An update. Clinical Neurophysiology: Official Journal of the International Federation of Clinical Neurophysiology, 122, 1686.

Rovo, Z., Matyas, F., Bartho, P., Slezia, A., Lecci, S., Pellegrini, C., Astori, S., David, C., Hangya, B., Luthi, A., & Acsady, L. (5 2014). Phasic, Nonsynaptic GABA-A Receptor-Mediated Inhibition Entrains Thalamocortical Oscillations. Journal of Neuroscience, 34, 7137–7147.

Russo, S., Sarasso, S., Puglisi, G. E., Palù, D. D., Pigorini, A., Casarotto, S., D’Ambrosio, S., Astolfi, A., Massimini, M., Rosanova, M., & Fecchio, M. (3 2022). TAAC - TMS Adaptable Auditory Control: A universal tool to mask TMS clicks. Journal of Neuroscience Methods, 370, 109491.

Salih, F., Khatami, R., Steinheimer, S., Hummel, O., Kühn, A., & Grosse, P. (6 2005). Inhibitory and excitatory intracortical circuits across the human sleep-wake cycle using paired-pulse transcranial magnetic stimulation. The Journal of Physiology, 565, 695–701.

Schaworonkow, N., Triesch, J., Ziemann, U., & Zrenner, C. (1 2019). EEG-triggered TMS reveals stronger brain state-dependent modulation of motor evoked potentials at weaker stimulation intensities. Brain Stimulation, 12, 110–118.

Sirota, A., Csicsvari, J., Buhl, D., & Buzsáki, G. (2 2003). Communication between neocortex and hippocampus during sleep in rodents. Proceedings of the National Academy of Sciences, 100, 2065–2069.

Somogyi, P., Katona, L., Klausberger, T., Lasztóczi, B., & Viney, T. J. (2 2014). Temporal redistribution of inhibition over neuronal subcellular domains underlies state-dependent rhythmic change of excitability in the hippocampus. Philosophical Transactions of the Royal Society of London. Series B, Biological Sciences, 369, 20120518.

Staresina, B. P., Bergmann, T. O., Bonnefond, M., van der Meij, R., Jensen, O., Deuker, L., Elger, C. E., Axmacher, N., & Fell, J. (11 2015). Hierarchical nesting of slow oscillations, spindles and ripples in the human hippocampus during sleep. Nature Neuroscience, 18, 1679–1686.

Stefanou, M.-I., Desideri, D., Belardinelli, P., Zrenner, C., & Ziemann, U. (12 2018). Phase Synchronicity of μ-Rhythm Determines Efficacy of Interhemispheric Communication Between Human Motor Cortices. The Journal of Neuroscience: The Official Journal of the Society for Neuroscience, 38, 10525–10534.

Szabo, G. G., Farrell, J. S., Dudok, B., Hou, W.-H., Ortiz, A. L., Varga, C., Moolchand, P., Gulsever, C. I., Gschwind, T., Dimidschstein, J., Capogna, M., & Soltesz, I. (6 2022). Ripple-selective GABAergic projection cells in the hippocampus. Neuron, 110, 1959–1977.e9.

Thankachan, S., Katsuki, F., McKenna, J. T., Yang, C., Shukla, C., Deisseroth, K., Uygun, D. S., Strecker, R. E., Brown, R. E., McNally, J. M., & Basheer, R. (12 2019). Thalamic Reticular Nucleus Parvalbumin Neurons Regulate Sleep Spindles and Electrophysiological Aspects of Schizophrenia in Mice. Scientific Reports, 9, 3607.

Thies, M., Zrenner, C., Ziemann, U., & Bergmann, T. O. (9 2018). Sensorimotor mu-alpha power is positively related to corticospinal excitability. Brain Stimulation, 11, 1119–1122.

Timofeev, I. (3 2001). Contribution of intrinsic and synaptic factors in the desynchronization of thalamic oscillatory activity. Thalamus & Related Systems, 1, 53–69.

Vallat, R., & Walker, M. P. (10 2021). An open-source, high-performance tool for automated sleep staging. eLife, 10. 10.7554/eLife.70092

Wischnewski, M., Haigh, Z. J., Shirinpour, S., Alekseichuk, I., & Opitz, A. (9 2022). The phase of sensorimotor mu and beta oscillations has the opposite effect on corticospinal excitability. Brain Stimulation, 15, 1093–1100.

Zrenner, C., Belardinelli, P., Ermolova, M., Gordon, P. C., Stenroos, M., Zrenner, B., & Ziemann, U. (9 2022). µ-rhythm phase from somatosensory but not motor cortex correlates with corticospinal excitability in EEG-triggered TMS. Journal of Neuroscience Methods, 379, 109662.

Zrenner, C., Desideri, D., Belardinelli, P., & Ziemann, U. (3 2018). Real-time EEG-defined excitability states determine efficacy of TMS-induced plasticity in human motor cortex. Brain Stimulation, 11, 374–389.

Zrenner, C., Galevska, D., Nieminen, J. O., Baur, D., Stefanou, M.-I., & Ziemann, U. (7 2020). The shaky ground truth of real-time phase estimation. NeuroImage, 214, 116761.

Zrenner, C., Kozák, G., Schaworonkow, N., Metsomaa, J., Baur, D., Vetter, D., Blumberger, D. M., Ziemann, U., & Belardinelli, P. (2023). Corticospinal excitability is highest at the early rising phase of sensorimotor µ-rhythm. NeuroImage, 266, 119805.

